# Pectin self-assembly and its disruption by water: Insights into plant cell wall mechanics

**DOI:** 10.1101/2021.12.03.471127

**Authors:** Jacob John, Debes Ray, Vinod K. Aswal, Abhijit P. Deshpande, Susy Varughese

## Abstract

Plant cell walls undergo multiple cycles of dehydration and rehydration during their life. Calcium crosslinked low methoxy pectin is a major constituent of plant cell walls. Understanding the dehydration-rehydration behavior of pectin gels may shed light on the water transport and mechanics of plant cells. In this work, we report the contributions of microstructure to the mechanics of pectin-Ca gels subjected to different extents of dehydration and subsequent rehydration. This is investigated using a pectin gel composition that forms ‘egg-box bundles’, a characteristic feature of the microstructure of low methoxy pectin-Ca gels. Large Amplitude Oscillatory Shear (LAOS) rheology along with Small Angle Neutron Scattering and Near Infrared (NIR) spectroscopy on pectin gels are used to elucidate the mechanical and microstructural changes during dehydration-rehydration cycles. As the extent of dehydration increase, the reswelling ability, strain-stiffening behavior and the yield strain decreases. These effects are more prominent at faster rates of dehydration and are not completely reversible upon rehydration to the initial undried state. Microstructural changes due to the aggregation of egg-box bundles and single chains and the associated changes in the water configurations lead to these irreversible changes.

## 1 Introduction

Water plays an important role in the mechanics of plant cell walls and thereby controls several crucial functions of plants such as the expansive growth [1] and stomatal functions [2]. Water transport within the apoplast is known to regulate stimuli responsive rapid movements in plants [3]. Several physical and chemical changes that can affect the viscoelastic properties, occur within the cell wall during hydration-dehydration cycles. Dehydration of Alfalfa stems are shown to result in severe distortion of shape of pectin-rich (non-lignified) cell walls due to physical changes in the polysaccharide network [4]. Distortion of shape and folding of cell walls are known to happen, which also act to preserve structural integrity during dehydration [5]. This is also known to be associated with the aggregation of the polysaccharide domains in the cell wall [6]. Compositional analysis has shown that the amount of extractable calcium crosslinked pectin decreased in the case of dehydrated cell walls indicating that dehydration results in the aggregation of the pectin chains due to non-covalent interactions [4]. The abundance of de-esterified (low methoxy) pectin and calcium crosslinking are associated with several important functions in plants such as the opening and closing of the stomata performed by the guard cells [7], the rehydration of pollen and germination [8] and organ initiation at apical meristem [9]. During dehydration, pectin is known to control the flexibility of the cell walls [10]. The essential regions are maintained flexible by creating an abundance of neutral sugar chains such as arabinose, which act as plasticizers in the polysaccharide network [10]. On the other hand, increased pectin-Ca crosslinks are induced in order to maintain the rigidity of certain non-essential cell walls [6]. The removal of water from the cell walls during desiccation is also known to alter the molecular structure and bonds among water molecules in addition to the changes in the polysaccharide organization [11].

One key feature of the microstructure of pectin-Ca gels is the parallel alignment of galacturonic acid chains with calcium ions at the center to form egg-box bundles. The extent of egg-box bundling is known to vary depending on the calcium ion content, extent of esterification, pectin concentration and the pectin chain conformation [12, 13], and leads to rheological behavior such as strain-stiffening and viscous dissipation. The removal of water by dehydration and its effects on the rheological behavior due to changes in the pectin-Ca gel microstructure are not well understood and are relevant for the mechanics of pectin-Ca gels as well as in the context of plant physiology. Hence, the effect of dehydration and rehydration on the microstructure of pectin-Ca gels and the associated changes on the rheological behavior are examined in this work.

Ca-crosslinked pectin being an integral part of the plant cell walls, we address the following questions in this work: what are the changes in the gel microstructure during dehydration and rehydration? How do these changes affect the mechanical response of the gel? Are the changes reversible during rehydration of the gel? Do the associated water molecules undergo changes in configuration due to microstructural changes? What is the role of Ca-mediated egg-box bundles during dehydration and rehydration? Large amplitude oscillatory shear (LAOS) rheological studies along with small angle neutron scattering (SANS) and near infrared (NIR) spectroscopy were carried out to answer these questions. Although there are many studies on the plant desiccation and rehydration behavior, to the best of our knowledge there are no studies on pectin-Ca gels to understand how calcium crosslinked pectin gels behave invitro.

## 2 Methods

### Materials

Pectin, from sunflower (*Helianthus annuus*), with galacturonic acid content 69%, degree of esterification of 33% and molecular weight (M_w_=32 kDa), used in this work was purchased from Krishna Pectins, Jalgaon, India. The FTIR spectra of the pectin used in this study was used for comparison to the chemical structure of the pectin found in plant cell walls (Figure S3, Supporting Information) [14]. CaCl_2_.2H_2_O was procured from Merck Specialties. Sodium chloride (NaCl 99% pure) was procured from Thermo Fisher Scientific. Deuterium Oxide (D_2_O, 99.9% atom D) required for the SANS experiments was procured from Sigma Aldrich.

### Drying and Swelling studies

Pectin-Ca gels with *R*=0.5 and pectin concentration of 1 wt% were prepared according to the procedure given in [12]. The dehydration of the gels were performed in an environment chamber (Memmert, CTC-256) and the gel samples were placed in a petri dish of 5 cm diameter, according to the procedure followed by Sekine et al., [15]. Effect of rate of dehydration was also studied by performing dehydration at different conditions such as, 25 °C and 50 % RH; 40 °C and 30 % RH; and 60 °C and 30 % RH, which corresponds to dehydration rates *k*=0.0033 g/h, *k*=0.0075 g/h, and *k*=0.0331 g/h respectively obtained from the slopes of the weight vs time curves during dehydration. Rehydration of the gels after dehydration was performed in deionized water at 25 °C. The swelling ratio, *Q*_*w*_, is obtained from,

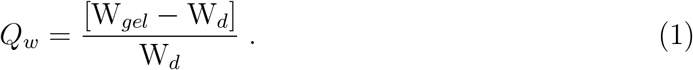

where, W_*gel*_ is the weight of the gel and W_*d*_ is the dry weight of the gel. *Q*_*w*_ is the weight ratio of the water to the polymer present in the gels and is used to describe the level of hydration in the gels.

### Rheological behavior

Large amplitude oscillatory shear (LAOS) rheology studies were carried out on pectin-Ca gels subjected to dehydration and rehydration. The experiments were carried out on Anton Paar, MCR-301, stress-controlled rheometer. Measurements were carried out using a cone and plate geometry of 50 mm diameter with a cone angle of 1.998°. In order to prevent the evaporation of water from the sample during the rheological measurements a solvent trap was used.

### NIR spectroscopy

The absorbance spectra for dehydrated and rehydrated gels were obtained using a Perkin-Elmer Lambda 950 UV-VIS-NIR spectrophotometer. The measurements were performed in the range of wavelength 200-2500 nm, with a step size of 1 nm at a temperature of 25 °C. The average from 8 consecutive measurements performed in reflectance mode is taken and the absorbance data is obtained as a function of wavelength.

### Small angle neutron scattering (SANS)

The neutron scattering measurements on dehydrated and rehydrated gels were performed at the Dhruva reactor, Bhabha Atomic Research Center, Mumbai, India. In order to enhance the contrast of the scattering, gels prepared in Deuterium Oxide D_2_O were used for the preparation of the dehydrated and rehydrated samples. Incident neutron beam with a mean wavelength (λ) of 5.2 °A and a resolution (*δ* λ/λ) of 10 % was used. A position sensitive *He*^3^ detector was used for measuring the scattered neutrons. Scattering vector, Q, ranged from 0.017 to 0.35 °A^-1^ (Q= 4*π*sin(*θ*/2)/*λ*, where *θ* is the scattering angle).

## 3 Results and Discussion

### Drying and rehydration behavior of the gels

Pectin-Ca gels (*R*=0.5), which is known to form egg-box bundles [12], are used in this study. The effects of extent of dehydration and the rate of dehydration on the rehydration behavior is shown in Figure 1 (a) in terms of the swelling ratio (*Q*_*w*_). During dehydration, the swelling ratio decreases from *Q*_*w*_=110.11, at its initial undried state to *Q*_*w*_ ≈0, at complete dehydration, for all rates of dehydration (*k*). The rehydration behavior of the gels after complete dehydration at different rates is shown in Figure 1 (b). The equilibrium value of *Q*_*w*_ of the rehydrated gels is lower compared to the initial *Q*_*w*_ (before dehydration), at all the dehydration rates studied, indicating the loss of ability of the gels to swell to their initial water content after complete dehydration. Also the equilibrium value of *Q*_*w*_ attained is lower when dehydration is performed at higher rates.

**Figure 1:**
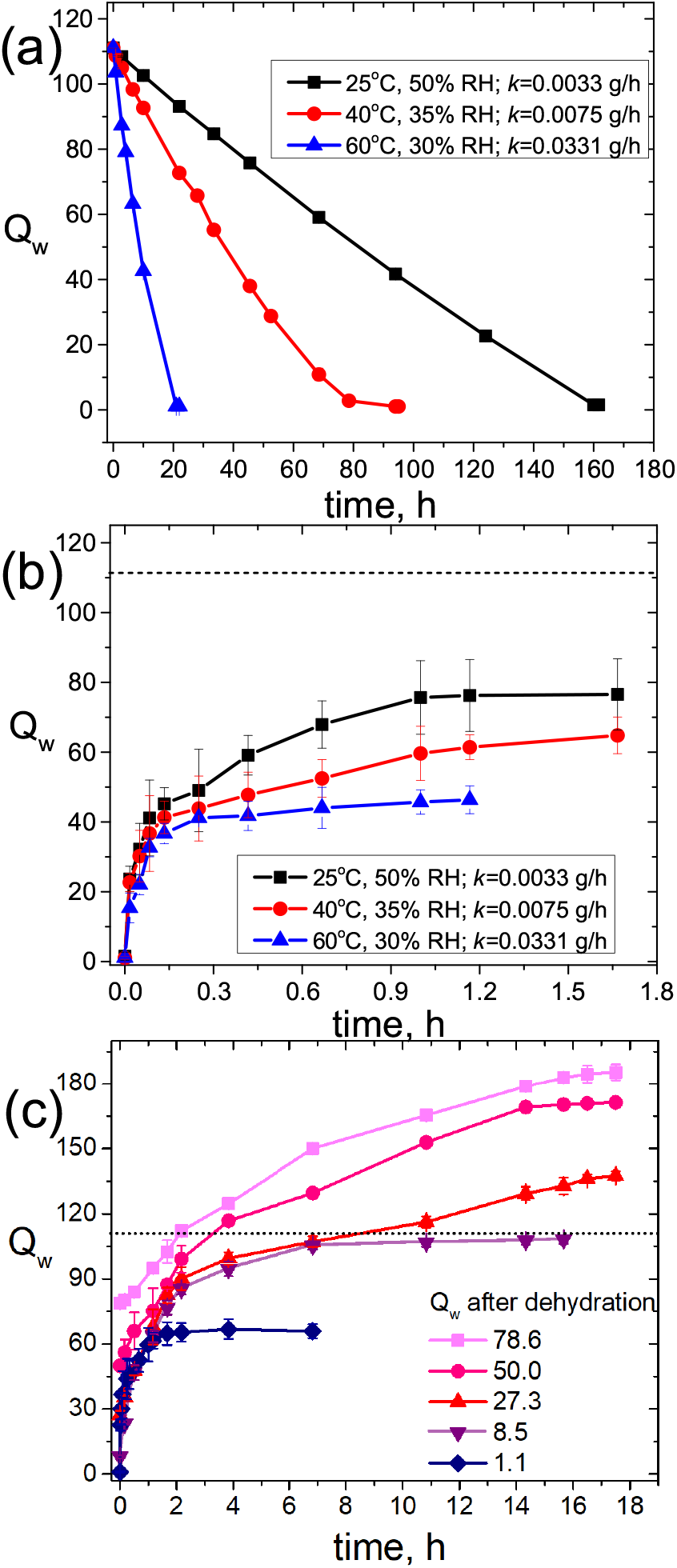
(a) Kinetics of dehydration of pectin-Ca gels (b) Kinetics of hydration following complete dehydration at different rates and (c) Kinetics of hydration following dehydration at k=0.0075 g/h to different extents. The dashed lines in (b) and (c) indicate the initial value of *Q*_*w*_ for the undried gels.

The incomplete reversal of the *Q*_*w*_ upon rehydration indicates irreversible microstructural changes induced during dehydration and this is more pronounced at higher drying rates. Figure 1 (c) shows the rehydrating behavior of the gels dehydrated to different water contents. The equilibrium water content attained during rehydration depends on the extent to which the gels were subjected to dehydration prior to the rehydration. Previous studies on dehydration and rehydration of non-resurrection type angiosperms have shown that survival of the plant is not possible if the water content reduces to 20-30% of that of the fully hydrated state[16]. In the case of bryophytes[17] and angiosperms[18], the ability to survive desiccation was found to reduce at faster dehydration rates. Pectin being the hydrophilic component of the plant cell walls, understanding its water storage behavior under various conditions is relevant here. To understand the possible contributions of calcium crosslinked pectin, the changes in the rheological behavior and the microstructure due to dehydration and rehydration are analyzed further.

### Mechanical response of dehydrated and rehydrated gels

The effect of dehydration and rehydration on the mechanical response of the gels is studied using LAOS rheology. Figure 2 (a) shows the effect of dehydration on the amplitude sweep response of the gel. When dehydrated from the initial state (*Q*_*w*_=110.11) to *Q*_*w*_=30.25, a three order increase in G*′* and a decrease in yield strain (*γ*_*c*_) are observed. For comparison, the rheological response of the undried gel with water content similar to that of the dehydrated gel (*Q*_*w*_=30.25) is also shown. Despite having similar water content, the dehydrated and the undried gels have large differences in the G*′* and *γ*_*c*_ values. These indicate that the microstructure and the crosslink density of the dehydrated and undried gels are different even at similar water contents. From the theory of rubber elasticity, the average crosslink density (*ν*) of the gels is known to be proportional to 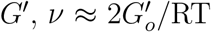 [19]. At similar *Q*_*w*_ values, the 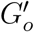 of the undried gels are approximately ten times higher than that of the dehydrated gels, indicating a higher crosslink density in the case of undried gels compared to the dehydrated gels. The dehydration at faster rates makes the gels yield at lower strains as can be observed from Figure 2 (b). The undried gels also undergo yielding at lower strains compared to the dehydrated gels.

**Figure 2:**
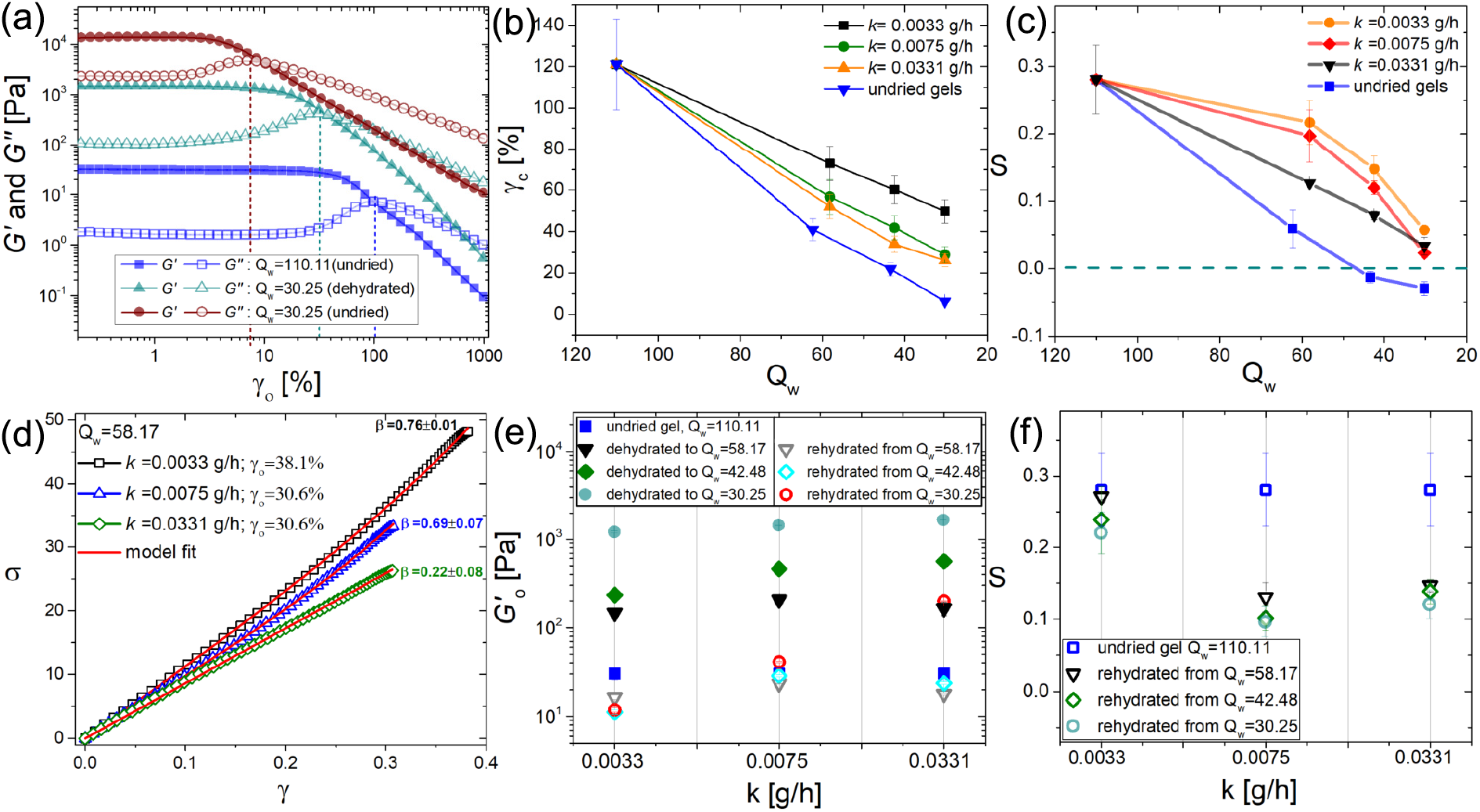
(a) Variation of *G*′ (closed symbols) and *G*^′′^ (open symbols) as a function of strain amplitude (*γ*_*o*_) for undried gel (*Q*_*w*_=110.11) and dehydrated gel (*Q*_*w*_=30.25). The behavior of undried gel with *Q*_*w*_=30.25 is also shown. (b) Variation of yield strain (*γ*_*c*_) as a function of *Q*_*w*_. The dotted lines in (a) indicates the yield strain (*γ*_*c*_). (c) Variation of strain-stiffening index (*S*) with the extent of dehydration. (d) Model fit to the stress-strain behavior from the first quarter of the Lissajous plots for pectin-Ca gels with *Q*_*w*_=58.17. (e) Variation of 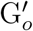 of the rehydrated gels as a function of dehydration rate. The dehydrated gels are represented using closed symbols and the corresponding rehydrated gels are represented using open symbols. (f) The variation of *S* for the rehydrated gels as a function of the dehydration rate.

The effect of dehydration is strongly evident in the strain-stiffening response during a cycle of oscillatory shear of the gels. Strain-stiffening index, ‘S’ (defined in Figure S1, Supporting information), decreases with decrease in *Q*_*w*_ and increasing dehydration rate, *k*. Interestingly, the undried gels have lower values of ‘S’ compared to that of the dehydrated gels at similar water contents. The undried gels have negative ‘S’ values at *Q*_*w*_ < 42, indicating a transition from strain-stiffening to strain-softening behavior. A phenomenological model developed by Dobrynin et al., is used for understanding the mechanism of strain-stiffening in the present case [20].

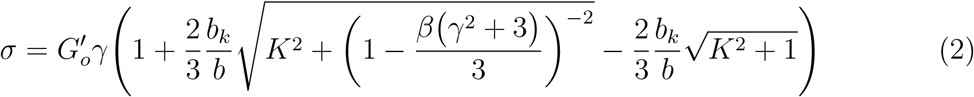

Here, 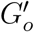 is the linear storage modulus, *b*_*k*_ is the Kuhn length of pectin molecules, *b* is the bond length between the monomers, *β* represents the chain elongation ratio, and *K* is the chain bending constant. Generally, *β* value close to 0.1 is obtained for networks with flexible chains, such as, rubber and ≈ 0.9 is obtained for semiflexible networks, such as, actin [21]. Figure 2 (d) shows stress-strain data in the first quarter of the Lissajous plots and its model fit using Equation 2 for the gels with *Q*_*w*_=58.17 obtained by dehydration from *Q*_*w*_=110.11, at different rates. This exercise was performed for the gels dehydrated at different rates of dehydration, to different *Q*_*w*_ values. The fitting parameter *β* has contributions from the underlying microstructure. With decrease in *Q*_*w*_ from 110.11 to 30.25, *β* decreases from ≈0.75 to ≈0.1 for all dehydration rates (Table S1, Supporting Information). This indicates that, with increase in dehydration, mostly the flexible single chains contribute to the stress response during shear. This could happen if the egg-box bundles and single chains form random shaped aggregates connected using single pectin chains which was confirmed using microstructural studies. This effect is observed to be more prominent at faster dehydration rates. Aggregation of polymer chains resulting in larger structures has been reported earlier in the case of alginate-Ca gels during dehydration [22]. Aggregation of polysaccharides and macromolecular chains due to dehydration is known to occur in plant cell walls [6] and synthetic polymer hydrogels respectively [15].

The effect of rehydration on the rheological behavior of the gels are shown in Figures 2 (e) and (f). The 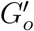 values for the rehydrated gels are closer to that of the undried gel, irrespective of the extent of dehydration, when dehydrated at slower rates (*k*=0.0033 g/h and *k*=0.0075 g/h). However at faster rates (*k*=0.0331 g/h), 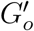 is dependent on the extent of dehydration. This indicates that, even though the dehydrated pectin-Ca gels with *R*=0.5 can be rehydrated to its initial water content, the underlying microstructure could be undergoing irreversible structural changes during dehydration. This is more evident in the changes in the strain-stiffening behavior of the rehydrated pectin-Ca gels. It can be observed from Figure 2 (f) that, rehydrated gels have a lower value of *S* compared to the undried gel. Also, the difference is higher in the case of the gels rehydrated after dehydrating at faster dehydration rates. The *β* values obtained from fitting the stress-strain data of the rehydrated gels using Equation 2 (Table S2, Supporting Information), also corroborates the observations made here. In the case of gels rehydrated after dehydrating at faster rate, the *β* values are lower. Microstructure and aggregation of the polymer chains are mediated by solvent configurations [15, 23] and therefore we investigate the H-bonding and clustering in different pectin-Ca gels to understand the changes in the microstructure during dehydration and rehydration.

### Changes in water configurations

The changes in the configuration of water molecules associated with pectin chains during the dehydration and rehydration of pectin-Ca gels was investigated using NIR spectroscopy, in the wavelength range of 1300-1700 nm, which corresponds to the overtones of the OH stretching peaks. Absorbance at specific wavelengths corresponding to distinct species of water configurations are termed as Water Matrix Coordinates (WAMACS) [24, 25] (Table S3, Supporting information). The relative value of the absorbance (A*′*) at different wavelengths corresponding to the WAMACS used for the construction of the aquagrams is obtained from,

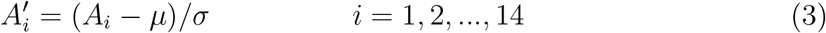

where *A*_*i*_. *µ* and *σ* are the absorbance value at wavelength *λ*, mean absorbance and standard deviation respectively. From Figure 3 (a), it can be observed that, with a decrease in *Q*_*w*_, there is an increase in the normalized absorbance at wavelengths, 1438, 1447, 1464, 1475, and 1492 nm. Simultaneously, a decrease in the normalized absorbance is observed at wavelengths 1343, 1358, 1371, 1395 and 1408 nm. These observed variations in the aquagram patterns indicate that dehydration result in a reduction in the amount of weakly hydrogen bonded water and protonated water clusters. This is accompanied by a simultaneous increase in the relative amounts of strongly hydrogen bonded water molecules. Water molecules in strongly H-bonded states can act as mediators that bind polymer molecules through H-bonding [26, 27]. An increase in the relative amount of water molecules having one, two and three H-bonds and less water clusters could be associated with the aggregation of pectin chains or egg-box bundles. These changes in water configurations support the analysis of the stress-strain data that suggests a loss in the semiflexible nature of the pectin chains due to the aggregation of chains and egg-box bundles.

**Figure 3:**
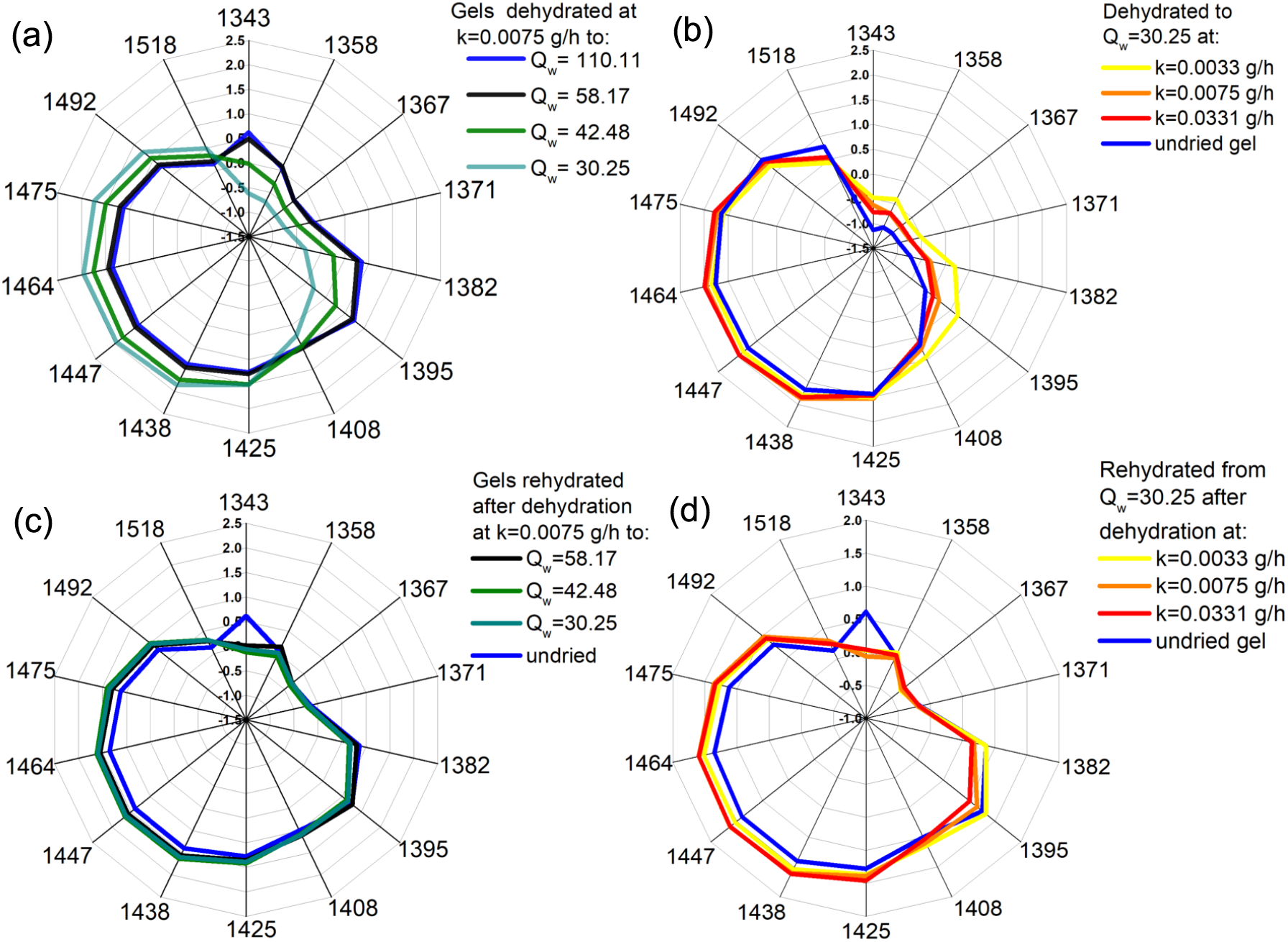
Aquagram patterns obtained from NIR spectroscopy of pectin-Ca gels (a) dehydrated at k=0.0075 g/h to different *Q*_*w*_ values. (b) dehydrated to *Q*_*w*_=30.25 at different dehydration rates (c) rehydrated to *Q*_*w*_=110.11 after dehydration at k=0.0075 g/h to different *Q*_*w*_ values. (d) rehydrated to *Q*_*w*_=110.11 after dehydration at different rates to *Q*_*w*_=30.25.

Further, Figure 3 (b) indicates that the changes in water configurations also depend on the rate of dehydration and are more significant at faster dehydration rates. Upon rehydration from different extents of dehydration, the water configurations do not reverse to its initial state, as is evident from the aquagram patterns of the rehydrated gels (Figure 3 (c)). The irreversibility of the changes in configurations of water molecules is more evident at faster dehydration rates (Figure 3(d)).

### Microstructural changes elucidated using SANS

Small angle neutron scattering (SANS) studies were carried out on dehydrated and rehydrated gels to examine changes in the microstructure of pectin-Ca gels. Figures 4 (a) and (b) show the scattering profiles for the gels, dehydrated to different extents at *k*=0.0033 g/h and *k*=0.0331 g/h respectively. A steep upturn in the intensity can be observed for *Q*_*w*_=30.25 at the lower dehydration rate (*k*=0.0033 g/h). This is more prominent in the case of the faster dehydration rate (*k*=0.0331 g/h) and at lower extent of dehydration. These changes generally indicate aggregation behavior in gels [28]. In the case of pectin-Ca gels this could be likely due to aggregation of the polymer chains and the egg-box bundles. Dehydration of alginate-Ca gels is known to result in aggregation of the junction zones and chains [22]. The microstructural changes resulting from aggregation are found to be gradual at slow dehydration rates and more drastic at faster dehydration rates. This indicates that when dehydration of the gels is performed at a faster rate, relatively larger structures are formed with corresponding smaller changes in the water content (Table S4, supporting information).

**Figure 4:**
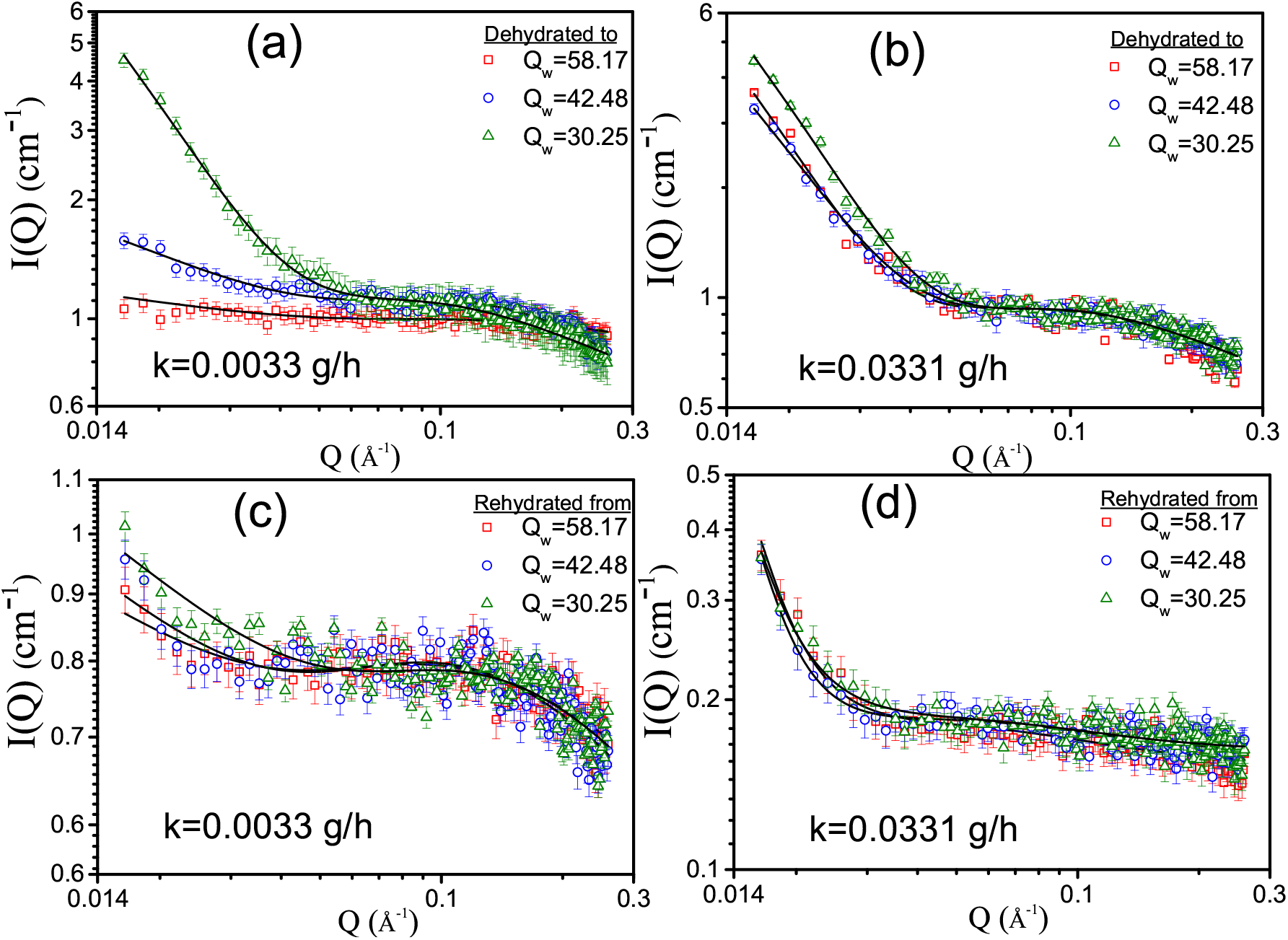
Scattering profiles for the pectin-Ca gel dehydrated to different extents at dehydration rates (a) *k*=0.0033 g/h and (b) *k*=0.0331 g/h and also for rehydrated gels after dehydration performed at (c) *k*=0.0033 g/h and (d) *k*=0.0331 g/h.

The analysis of the scattering data was performed using a model that combines the contributions of scattering of single chains and the fractal nature of the gel network which is made up of rod-like egg-box bundles. Hence, the scattering intensity (*I*(*Q*)), was fitted using the form factor given below to find the size of the aggregates (*R*_*g*_),

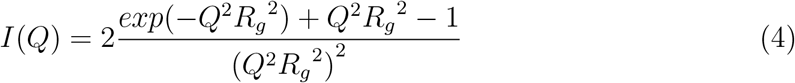

The shape of the aggregates is determined by estimating the fractal dimension (*d*_*f*_) of the aggregates, using a power law relation, 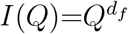. As evident from the qualitative variation of the scattering pattern with dehydration, the aggregate size (*R*_*g*_) increases with dehydration (Table S4, Supporting Information). The X-ray diffractogram of the dehydrated gel also indicates an increase in the size of the aggregates (Figure S2, Supporting Information). The increase in the fractal dimension from *∼*1 to *∼*1.8 indicates a transition in shape of the aggregates from rod-like to a random coiled aggregate. Similar variation was observed and was found to be more drastic at faster dehydration rates.

Upon rehydration, the absolute scattering intensity *I*(*Q*) reduces as shown in Figures 4 (c) and (d), due to the reduction in polymer concentration due to swelling. Interestingly, the steep upturn observed at low *Q* values in the case of the dehydrated gels is also present in the case of the rehydrated gels, indicating the persistence of larger aggregates formed during dehydration after rehydration. This is more prominent at faster dehydration rates and higher extents of dehydration. Analysis of the scattering data for the rehydrated gels shows that the fractal dimension changes from *∼*1.8 to *∼*1 in the case of the slower dehydration rates and remains *∼*2 in the case of faster dehydration rates (Table S5, Supporting Information). The observations from rheology, aquagrams and SANS are summerized schematically in Figure 5. The change in microstructure is consistent with the diminishing strain-stiffening behavior observed and the analysis of the stress-strain data using Equation 2. The increasing extent of aggregation implies major contribution of the single chains connecting the aggregates, deforming under shear and correspondingly reduced contributions from egg-box bundles. Variation in the water configurations near the aggregates is also shown in the schematic.

**Figure 5:**
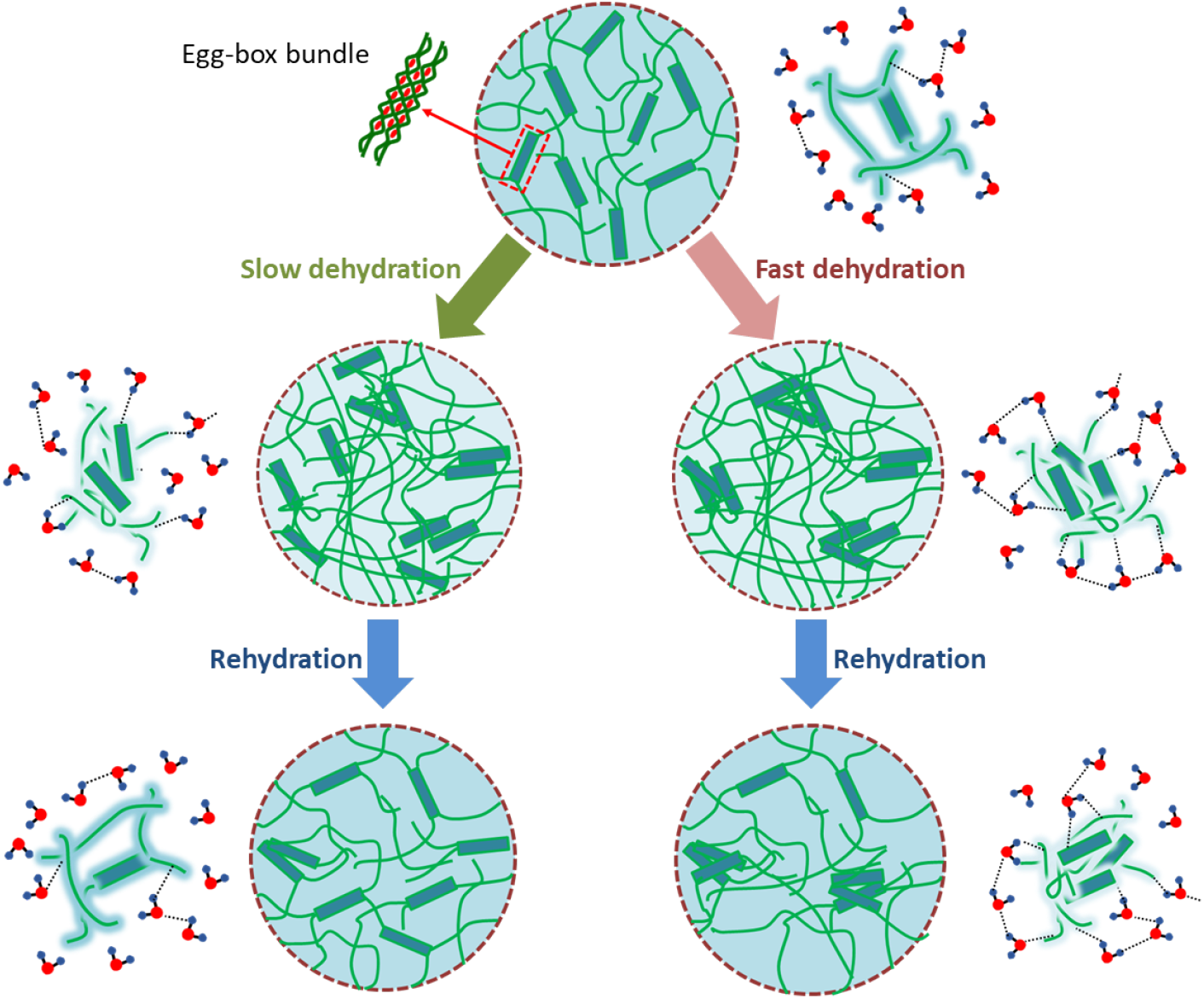
Schematic representation of the microstructural changes observed using rheol-ogy, aqugrams and SANS.

## 4 Conclusion

This work shows the mechanisms through which low methoxy pectin in the Ca-crosslinked form could influence the dehydration and rehydration behavior in plant cell walls. Dehydration of pectin-Ca gels decreases the hydrating ability and diminishes its strain-stiffening behavior due to the induced microstructural changes accompanied by changes in water configurations. The mentioned effects and its reversibilty upon rehydration are dependent on the rate of dehydration and the extent of dehydration. Microstructural changes such as the aggregation of egg-box bundles and single chains found in the gels could determine the dehydration rate dependent rehydration ability observed in plants [29]. The diminishing strain-stiffening behavior observed during dehydration of pectin-Ca gels could affect the functions of the pectin rich tissues such as the shoot apical meristem [30] and the guard cells of the stomata [31], where strain-stiffening is important. Plants take up water along with salts and minerals which may also play important roles in the dehydration and rehydration behavior in plants. Therefore the role of active restoration of salts like calcium and other minerals needs to be investigated along with water in order to obtain a complete picture.

## References

[1] Daniel J. Cosgrove. Plant cell wall extensibility: connecting plant cell growth with cell wall structure, mechanics, and the action of wall-modifying enzymes. Journal of Experimental Botany, 67(2):463–476, 11 2015.

[2] Thomas N. Buckley. How do stomata respond to water status? New Phytologist, 224(1):21–36, 2019.

[3] Jacques Dumais and Yöel Forterre. “vegetable dynamicks”: The role of water in plant movements. Annual Review of Fluid Mechanics, 44(1):453–478, 2012.

[4] M. Hamm, P. Debeire, B. Monties, and B. Chabbert. Changes in the cell wall network during the thermal dehydration of alfalfa stems. Journal of Agricultural and Food Chemistry, 50(7):1897–1903, 2002.

[5] Mary Alice Webb and Howard J. Arnott. Cell wall conformation in dry seed sub relation to the preservation of structural integrity during desiccation. American Journal of Botany, 69(10):1657–1668, 1982.

[6] John P. Moore, Mäite Vicré-Gibouin, Jill M. Farrant, and Azeddine Driouich. Adaptations of higher plant cell walls to water loss: drought vs desiccation. Physiologia Plantarum, 134(2):237–245, 2008.

[7] Sam Amsbury, Lee Hunt, Nagat Elhaddad, Alice Baillie, Marjorie Lundgren, Yves Verhertbruggen, Henrik V. Scheller, J. Paul Knox, Andrew J. Fleming, and Julie E. Gray. Stomatal function requires pectin de-methyl-esterification of the guard cell wall. Current Biology, 26(21):2899–2906, 2016.

[8] Fan Teng-Fei, Park Soohyun, Shi Qian, Zhang Xingyu, Liu Qimin, Song Yoohyun, Chin Hokyun, Ibrahim Mohammed Shahrudin Bin, Mokrzecka Natalia, Yang Yun, Li Hua, Song Juha, Suresh Subra, and Cho Nam-Joon. Transformation of hard pollen into soft matter. Nature Communications, 11(2041-1723), 2020.

[9] Alexis Peaucelle, Siobhan A. Braybrook, Laurent Le Guillou, Emeric Bron, Cris Kuhlemeier, and Herman Höfte. Pectin-induced changes in cell wall mechanics underlie organ initiation in arabidopsis. Current Biology, 21(20):1720–1726, 2011.

[10] John P. Moore, Jill M. Farrant, and Azeddine Driouich. A role for pectin-associated arabinans in maintaining the flexibility of the plant cell wall during water deficit stress. Plant Signaling & Behavior, 3(2):102–104, 2008.

[11] Shinichiro Kuroki, Roumiana Tsenkova, Daniela Moyankova, Jelena Muncan, Hiroyuki Morita, Stefka Atanassova, and Dimitar Djilianov. Water molecular structure underpins extreme desiccation tolerance of the resurrection plant haberlea rhodopensis. Scientific Reports, 9(3049), 2019.

[12] Jacob John, Debes Ray, Vinod K. Aswal, Abhijit P. Deshpande, and Susy Varughese. Dissipation and strain-stiffening behavior of pectin–ca gels under laos. Soft Matter, 15:6852–6866, 2019.

[13] Erich Schuster, Aurelie Cucheval, Leif Lundin, and Martin A. K. Williams. Using saxs to reveal the degree of bundling in the polysaccharide junction zones of microrheologically distinct pectin gels. Biomacromolecules, 12(7):2583–2590, 2011.

[14] Maria H.G. Canteri, Catherine M.G.C. Renard, Carine Le Bourvellec, and Sylvie Bureau. Atr-ftir spectroscopy to determine cell wall composition: Application on a large diversity of fruits and vegetables. Carbohydrate Polymers, 212:186–196, 2019.

[15] Yurina Sekine and Tomoko Ikeda-Fukazawa. Structural changes of water in a hydrogel during dehydration. The Journal of Chemical Physics, 130(3):034501, 2009.

[16] Melvin J. Oliver, John C. Cushman, and Karen L. Koster. Dehydration Tolerance in Plants, pages 3–24. Humana Press, Totowa, NJ, 2010.

[17] Influence of dehydration rate on cell sucrose and water relations parameters in an inducible desiccation tolerant aquatic bryophyte. Environmental and Experimental Botany, 120:18 –22, 2015.

[18] Jill M. Farrant, Keren Cooper, Lynette A. Kruger, and Heather W. Sherwin. The effect of drying rate on the survival of three desiccation-tolerant angiosperm species. Annals of Botany, 84(3):371–379, 1999.

[19] L.R.G. Treloar. pages 71–74. Oxford University Press, London, 1975.

[20] Jan-Michael Y. Carrillo, Fred C. MacKintosh, and Andrey V. Dobrynin. Nonlinear elasticity: From single chain to networks and gels. Macromolecules, 46(9):3679–3692, 2013.

[21] Andrey V. Dobrynin and Jan-Michael Y. Carrillo. Universality in nonlinear elasticity of biological and polymeric networks and gels. Macromolecules, 44(1):140–146, 2011.

[22] Liangbin Li, Yapeng Fang, Rob Vreeker, Ingrid Appelqvist, and Eduardo Mendes. Reexamining the egg-box model in calciumalginate gels with x-ray diffraction. Biomacromolecules, 8(2):464–468, 2007.

[23] Kushi Kudo, Junichi Ishida, Gika Syuu, Yurina Sekine, and Tomoko Ikeda-Fukazawa. Structural changes of water in poly(vinyl alcohol) hydrogel during dehydration. The Journal of Chemical Physics, 140(4):044909, 2014.

[24] Li Ma, Xiaoyu Cui, Wensheng Cai, and Xueguang Shao. Understanding the function of water during the gelation of globular proteins by temperature-dependent near infrared spectroscopy. Phys. Chem. Chem. Phys., 20:20132–20140, 2018.

[25] Kodzue Kinoshita, Mari Miyazaki, Hiroyuki Morita, Maria Vassileva, Chunxiang Tang, Desheng Li, Osamu Ishikawa, Hiroshi Kusunoki, and Roumiana Tsenkova. Spectral pattern of urinary water as a biomarker of estrus in the giant panda. Scientific Reports, 2(856), 2012.

[26] Eri Chatani, Yutaro Tsuchisaka, Yuki Masuda, and Roumiana Tsenkova. Water molecular system dynamics associated with amyloidogenic nucleation as revealed by real time near infrared spectroscopy and aquaphotomics. PLOS ONE, 9:1–10, 07 2014.

[27] RN Tsenkova, IK Iordanova, K Toyoda, and DR Brown. Prion protein fate governed by metal binding. Biochemical and Biophysical Research Communications, 325(3):1005–1012, 2004.

[28] Brett D. Ermi and Eric J. Amis. Domain structures in low ionic strength polyelectrolyte solutions. Macromolecules, 31(21):7378–7384, 1998.

[29] Yongheng Liang and Wendell Q. Sun. Rate of dehydration and cumulative desiccation stress interacted to modulate desiccation tolerance of recalcitrant cocoa and ginkgo embryonic tissues. Plant Physiology, 128(4):1323–1331, 2002.

[30] Daniel Kierzkowski, Naomi Nakayama, Anne-Lise Routier-Kierzkowska, Alain Weber, Emmanuelle Bayer, Martine Schorderet, Didier Reinhardt, Cris Kuhlemeier, and Richard S. Smith. Elastic domains regulate growth and organogenesis in the plant shoot apical meristem. Science, 335(6072):1096–1099, 2012.

[31] Ross Carter, Hugh Woolfenden, Alice Baillie, Sam Amsbury, Sarah Carroll, Eleanor Healicon, Spyros Sovatzoglou, Sioban Braybrook, Julie E. Gray, Jamie Hobbs, Richard J. Morris, and Andrew J. Fleming. Stomatal opening involves polar, not radial, stiffening of guard cells. Current Biology, 27(19):2974 – 2983.e2, 2017.

